# Long distance runners in the marine realm: New insights into genetic diversity, kin relationships and social fidelity of Indian Ocean male sperm whales

**DOI:** 10.1101/2021.04.23.440733

**Authors:** Justine Girardet, Francois Sarano, Gaëtan Richard, Paul Tixier, Christophe Guinet, Alana Alexander, Véronique Sarano, Hugues Vitry, Axel Preud’homme, René Heuzey, Ana M. Garcia-Cegarra, Olivier Adam, Bénédicte Madon, Jean-Luc Jung

## Abstract

**Background:** Adult male sperm whales (*Physeter macrocephalus*) are long distance runners of the marine realm, feeding in high latitudes and mating in tropical and subtropical waters where stable social groups of females and immatures live. Several areas of uncertainty still limit our understanding of their social and breeding behaviour, in particular concerning the potential existence of geographical and/or social fidelities.

In this study, using underwater observation and sloughed-skin sampling, we looked for male social fidelity to a specific matrilineal sperm whale group near Mauritius. In addition, we captured a wider picture of kin relationships and genetic diversity of male sperm whales in the Indian Ocean thanks to biopsies of eight unique individuals taken in a feeding ground near the Kerguelen and Crozet Archipelagos (Southern Indian Ocean).

**Results:** Twenty-six adult male sperm whales, of which 13 were sampled, were identified when socializing with adult females and immatures off Mauritius. Long-term underwater observation recorded several noteworthy social interactions between adult males and adult females and/or immatures. We identified seven possible male recaptures over different years (three by direct observation, and four at the gametic level), which supports a certain level of male social fidelity. Several first- and second-degree kin relationships were highlighted between members of the social unit and adult males, confirming that some of the adult males observed in Mauritian waters are reproductive. Male social philopatry to their natal group can be excluded, as none of the males sampled shared the haplotype characteristic of the matrilineal social group. Mitochondrial DNA control region haplotype and nucleotide diversities calculated over the 21 total male sperm whales sampled were similar to values found by others in the Indian Ocean.

**Conclusions:** Our study strongly supports the existence of some levels of male sperm whale social fidelity, not directed to their social group of birth, in the Indian Ocean. Males sampled in breeding and feeding grounds are linked by kin relationships. Our results support a model of male mediated gene flow occurring at the level of the whole Indian Ocean, likely interconnected with large-scale geographical fidelity to ocean basin, and a small-scale social fidelity to matrilineal social groups.

## Background

Sexual dimorphism, defined as differences in external appearance or other characteristics between the two sexes of a species [1], is widespread among animals, and especially in vertebrates [2]. Sexual dimorphism can be behavioural, morphological and/or concern life history. Marked sexual dimorphism is present in several marine mammal species [1]. Morphological differences are obvious, for example, in elephant seals (*Mirounga angustirostris* and *M. leonina*), males being up to ten times larger than females [3] and in narwhals (*Monodon monoceros*) where males possess a tusk [4]. Other species display marked sexual segregation in geographical distribution, such as the Indo-Pacific bottlenose dolphins (*Tursiops aduncus*, [5, 6], or exhibit differences in their feeding ecology such as the resident fish-eating population of killer whales of the northeastern Pacific Ocean (*Orcinus orca*, [7].

Sperm whales certainly display some of the most striking sexual dimorphism among cetaceans, both in terms of body size with adult males growing up to 18m long and a weight of 45t, while females usually remain around 11m long for 13t [8, 9]; but also in terms of feeding ecology, geographical distribution and social organization [10–13]. Male and female sperm whales live in societies that are strongly geographically segregated post-maturity (e.g. [14–19]. Adult females form social units with immatures, stable over time and found all year round in warm waters at low latitudes [11, 20, 21]. In contrast, males disperse from their natal group after 6-8 years old, before their sexual maturity, and move poleward to areas abundant in food [10]. After their twenties, they make periodic forays to warmer waters for mating, with no known clear frequency, seasonal agendas nor migration routes [8]. Although we know that adult male sperm whales can travel thousands of kilometres across ocean basins [22, 23], no recurrent migration routes between feeding and breeding areas have so far been identified [9].

In cold waters, non-breeding adult males can be encountered alone or in small groups called “bachelor groups”, groups of tens individuals of about the same age (e.g. [24–26]. They may become more and more solitary as they age [8]. In northern Norway, Nova Scotia (Canada) and Kaikoura (New Zealand) feeding grounds, no noticeable social interaction between adult males were observed when foraging [24, 27]. Yet, some recent studies show that males can form long-term associations [13] and have fluid and unstructured social interactions that allow the social transmission of depredation techniques in the Gulf of Alaska [28] or permit coordinated anti-predator responses [29]. Long-term photo-identification studies around Crozet and Kerguelen archipelagos (Crozet/Kerguelen, Southern Indian Ocean), in the Bleik Canyon (northern Norway) and in the Nemuro Strait (northern Japan) indicate that adult males exhibit site fidelity at local scales [19, 30, 31].

In the low latitudes, the social interactions of adult male sperm whales with stable social groups of females and immatures and adult male movement patterns in breeding grounds remain poorly known. Adult males may temporarily join social units to breed and stay in the same area for periods estimated from a few hours to a few days off the Galapagos Islands [32] to few weeks in the West Indies [33]. During this period, large males roam around, apparently avoiding one another while visiting groups of females [9] and having limited social interactions with members of the social units (adult females and/or immatures, [33]. The existence of geographical and/or social fidelity is questioned in males, however fidelity of adult males to the ocean of their birth (i.e. a large geographical scale natal philopatry) has been suggested by whaling reports [8]. Using genetic assignment, Mesnick et al. [34] highlighted that, in the North Pacific, a higher-than randomly-expected proportion of males returned to their population of origin to mate. Males sharing first order kinships (mostly full siblings) have also been identified in the Azores and in the Chagos Archipelago [35, 36]. Photo-identification recaptures of a same male over several years in the same study area occurred in different breeding grounds of the Atlantic (in the Azores and the West Indies, [33, 37] and of the Pacific (The Galapagos, [14]), where they may socialise with different social groups of the same vocal clan [38]. Gero et al. [33] suggested that male fidelity to breeding sites might occur, based on the identification of the same male spanning a period of ten years and the observation of a gathering of dozens of females and immatures around a male.

Altogether, these results suggest that some levels of geographical and social fidelity could exist in male sperm whales. This hypothesis requires more evidence to be confirmed, however long-term monitoring of adult male sperm whales is difficult. Few studies have included males in analyses when studying female social groups (e.g. [32, 33, 35, 38]), and this scarcity of data prevents clear conclusions concerning male sperm whale movement patterns and social fidelity being drawn.

In the Indian Ocean breeding grounds, sperm whales have been less studied than in the Pacific and the Atlantic. Several social groups have been observed [11, 39, 40], and photoidentification campaigns and satellite tracks confirmed that sperm whales are common near the Mauritius and La Reunion Islands [40–42]. The predominant matrilineality of a particular social group, the “Irène’s group” has been recently demonstrated near Mauritius [21]. But except for some photo-identified individuals [40], male sperm whales encountered within the breeding grounds of the Indian Ocean are very poorly known. More knowledge comes from the feeding grounds of the Indian Ocean, and in particular from Crozet/Kerguelen [19, 43, 44], however the movement patterns between feeding and breeding grounds are not known.

In this study, we investigated the spatial and social fidelity of adult male sperm whales in the Indian Ocean. Using nine years of monitoring sperm whale social groups off Mauritius paired with genetic information collected on individuals from both this area and the Crozet/Kerguelen region, our aims were to: i) assess the association patterns and genetic relatedness of adult males with the members of a resident social group they associate with; ii) determine the extent of genetic relatedness across adult males, and, iii) analyse possible social and geographical fidelity of adult male sperm whales, including whether they show fidelity to their natal social group.

## Results

### 2011-2020 assessment of adult male sperm whale observations off Mauritius

A total of 26 adult males were identified based on their body length by underwater observations between 2011 and 2020 off Mauritius (Table S3). Males were observed in 2011, 2013 and yearly since 2015 when the observation effort significantly increased [40]. Since then, adult male sperm whales were sighted each year with a maximum of 10 different individuals observed in 2019. Adult males were observed during a total of 59d over the 2015-2020 period with a maximum of 29d in 2019 (Table S3). Observations of adult males occurred most of the year with at least one male seen each month from February to December. Over the 2015-2020 period of observations, April was the month with the highest rate of identification (seven males). Almost half of the males were identified on at least two different days within or between years (n=11), 15 were seen only once. When multiple sightings of the same male occurred during a given year, the longest span between the first and the last sightings was 47d (Léonard and Jason in 2019), with a mean of 8.25d (range = 1d-47d) (Table S3). Three males were positively identified over multiple years: Jonas, sighted in 2018 and 2019; Navin, sighted in 2015 and in 2018; and Hugues, sighted in 2013 and again 6 years later in 2019 (Table S3).

### Observation of particular social interactions between adult male sperm whales and members of the Irène’s group

Different socializing behaviours were observed between adult females and/or immatures of the Irene’s group. Figure 1 shows an example of an adult male (Reza) surrounded by an adult female and seven immatures (five males and two females) of the Irène’s group. This kind of socializing behaviour between an adult male and several immatures is not uncommon since it was observed and filmed 16 times in 2019, and involved 5 different adult males: Daniel, Reza, Léonard, Jason, Jonas.

**Figure 1.**
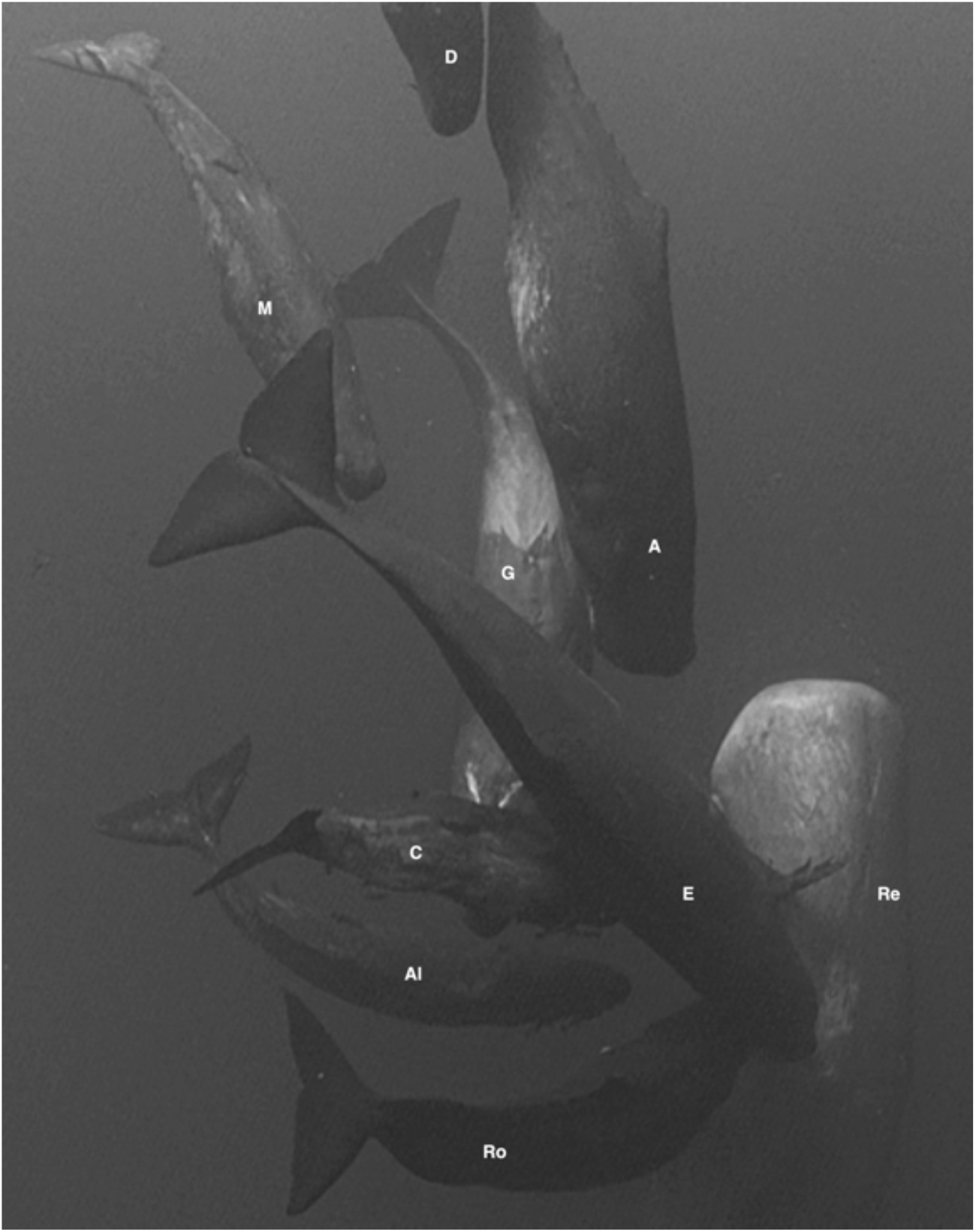
Social interactions between an adult male (Reza, **Re**), an adult female (Germine, **G**) and different immatures of the Irène’s group: Eliot (**E**) 8yrs-old; Arthur (**A**) and Roméo (**Ro**) 6yrs-old; Ali (**Al**) Daren (**D**) and Chesna (**C**) 1yr-old, and Miss Toutou (**M**) 3yrs-old.

The arrival of Jonas and Aman in July 2018 was also a particularly interesting event: this arrival initiated a large gathering of females and immatures of different social units. At least 60 females and immatures were observed at this time (MMCO, Field report of the July 18, 2018). Social interactions (e.g., *swimming together*) were also observed between adult males present in Mauritian waters at the same time. The most striking example of these social interactions was that of Jason and Léonard. Throughout their presence, from April 23, 2019 to June 8, 2019, they were observed together at each observation (n=11) (Table S3).

### Genetic analysis

A total of 132 sloughed skin samples were collected between 2017 and 2020 (Table S2). They were assigned in the field to 41 different sperm whales, i.e. to 18 adult females and 10 immatures (Table S2 and Sarano F. et al. 2021) and to 13 adult males (Table S2). Mitochondrial and nuclear loci were amplified, allowing an analysis of variation over 638 bp of the MCR (Genbank references: MK907146-MK907148, MK907159, MK907163, MK907172, MW854724-MW854731 and MW929445-MW929452) and at 16 polymorphic microsatellite loci (Table S1). The Identity Analysis based on microsatellite polymorphisms performed in CERVUS identified thirteen genetically distinct individuals from Mauritius corresponding to the 13 adult males identified in the field (all pID <2.45e^−12^). All genotypes assigned to the same individual had between 87.5% and 100% identity, and the differences were all consistent with allelic drop out. Mitochondrial haplotypes were all 100% identical between samples of the same individual. Only three skin samples had to be reassigned to another sperm whale than the one identified in the field after a *posteriori* careful examination of video recordings (see Table S2 and Sarano et al. 2021 for more explanation). Nine samples were taken off Crozet/Kerguelen, among which 8 genetic individuals were identified, Bio_Cro_2011_1 and Bio_Cro_2017 corresponding to the same individual (pID = 2.6e^−23^). Six MCR haplotypes were detected among the thirteen adult male sperm whales sampled off Mauritius (*H*=0.72, π=0.00265). Five different MCR haplotypes were identified in the eight male sperm whales sampled in Crozet/Kerguelen (*H*=0.78, π=0.00274). Mitochondrial *Φ_ST_* calculated between males sampled near Mauritius and those sampled in Crozet/Kerguelen was significant (*Φ_ST_*=0.136, p=0.037), and the *F_ST_* value was just above the significant value fixed to 5% (*F_ST_*=0.125, p=0.055).

### Genetic relationships between Irène’s social unit members and adult males sampled off Mauritius

In this study, the mitochondrial haplotype names correspond to the geographical places they came from (M: Mauritius, C: Crozet, K: Kerguelen). The correspondence with the haplotypes defined by Alexander et al. (2016) is presented in Table S4 and Figure 2. One adult male harboured the SW_M1 haplotype, corresponding to haplotype C of [36], characteristic of the Irène’s group [21]. Two others had the same haplotype (SW_MCK1) as Claire, the sole adult female of the Irene’s social group with a different MCR haplotype [21], corresponding to the haplotype N.001.001 mainly found in the Seychelles, in the Coco Islands and in the south west Australia by [36]. Another adult male had the haplotype MCK2 (differing from SW_MCK1 at position 609, table S4), one had the haplotype SW_M3 corresponding to the haplotype KK found almost exclusively in the Indian Ocean off Sri Lanka [36] and off Albany in Australia [45]. Seven males shared the haplotype SW_MC, identical to the haplotype A.001.001, common in the Indian Ocean (Figure 2). The last male possessed a new haplotype, SW_M2, not found previously anywhere else.

**Figure 2.**
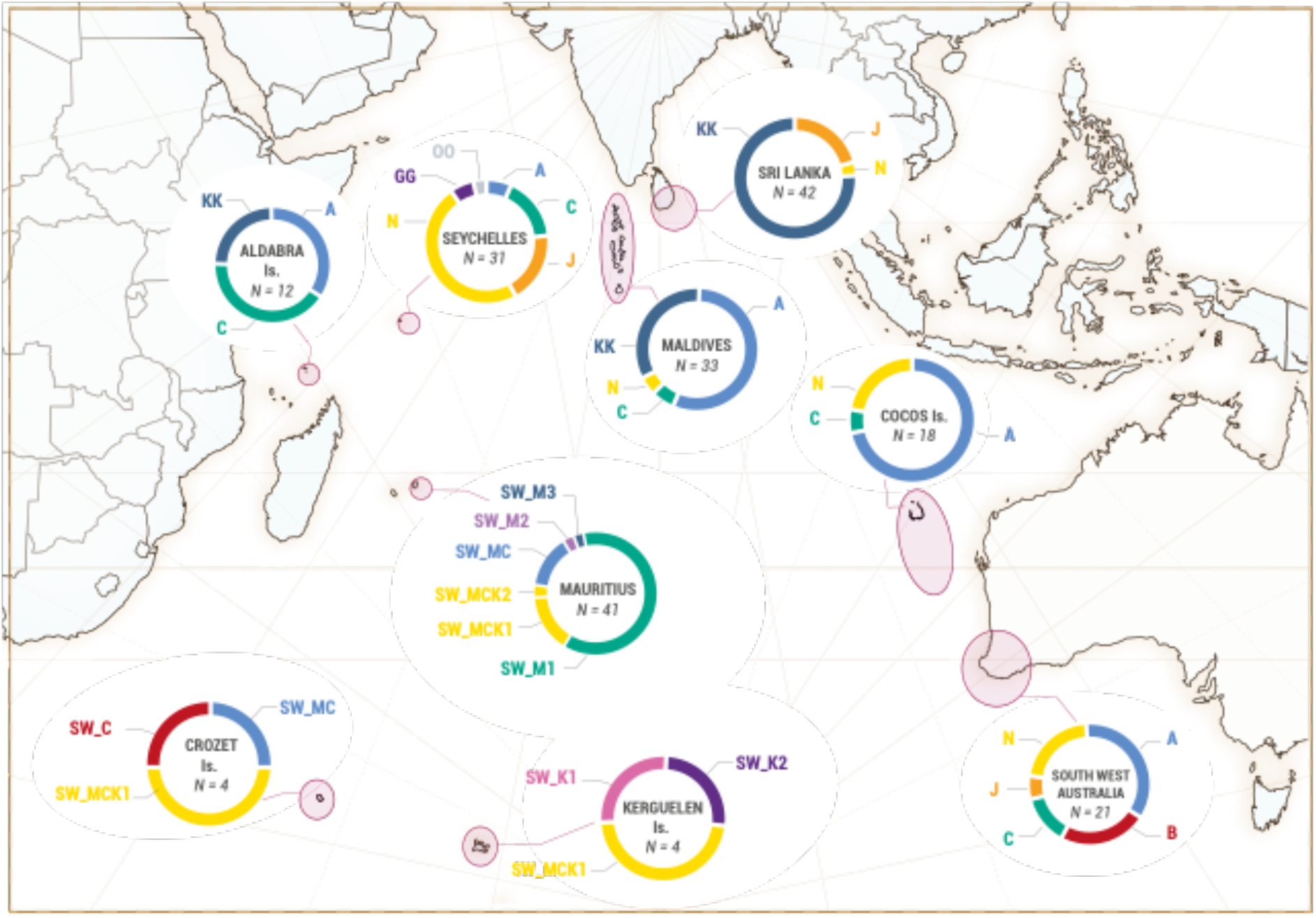
**Geographical repartition of the mitochondrial haplotypes in the Indian Ocean** determined by Alexander et al. 2016 (haplotypes named with one or two letters) and haplotypes determined during this study (haplotypes names starting by SW). A same colour indicates corresponding haplotypes (602bp in common). N: number of sperm whales for each diagram.

Kinship analysis revealed two first- and 20 second-degree kin relationships (11 with adult females, 9 with immatures) between the 13 adult males sampled in Mauritius and members of the Irène’s group (Figure 3 and Table S5). One adult male, Jonas, was identified as the father of Daren, a young male born in 2018; and a second adult male, Noé, was identified as the father of Lana, a young female born in 2019 (Figure 3, Table S5). All but three adult males presented at least one second-degree relationship with members of the Irène’s group with a maximum of four (Josuah and Léonard) (Figure 3).

**Figure 3:**
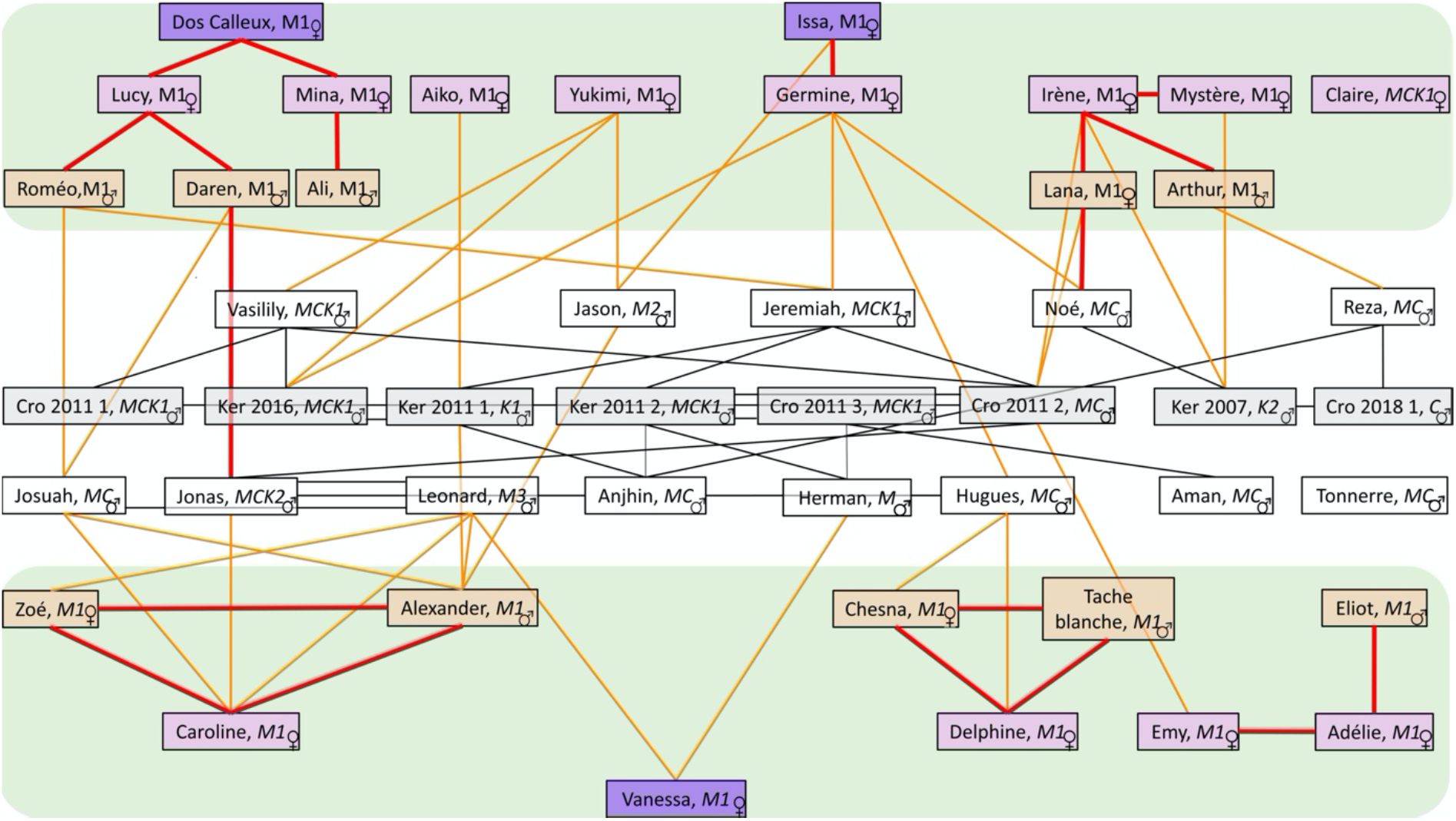
Schematic representation of the kin relationships between all the members of the Irène’s group and the adult males sampled off Mauritius (n=13) and in the Sub-Antarctic waters of the South of the Indian Ocean (n=8). First-degree (red lines) and second-degree (black lines between two adult males and orange lines between an adult male and a member of Irène’s group) relationships between the different sperm whales are represented (second degree between members of the Irène’s social group are not represented for the sake of clarity, see [21] for these relationships). The name, sex, and mitochondrial haplotypes (listed in Table S4) are indicated for each individual. Adult females are represented in purple (dark for older individuals, as estimated in the field, and light purple for the others), young sperm whales within the Irène’s social group in orange, and adult males are in white (males from Mauritius) and in light grey (males from Crozet/Kerguelen – sampling locations designated by “Cro” and “Ker”, respectively). The two green boxes represent two social subgroups identified within the Irène’s social group (Sarano et al. 2021). As stated in Sarano et al. (2021), this diagram was constructed to be consistent with the analyses conducted. Although we performed different analyses that produced similar results, uncertainty exists in the relatedness estimate calculations, which might influence some of these relationships.

Four possible full sibling relationships (same mother and father) have also been discovered in the Irène’s group (two between immatures and two between adult females).

### Large geographic scale kin relationships in the Indian Ocean

Two haplotypes (SW_MCK1 and SW_MC) were found both in Crozet/Kerguelen and in Mauritian males. SW_MCK1, shared by four sub-Antarctic sperm whales (two sampled in Crozet and two in Kerguelen) was the most frequent. The haplotype SW_MC was found in one sperm whale from Crozet (Figure 2). Three other haplotypes were found in the Crozet/Kerguelen samples that were not observed among males sampled off Mauritius: SW_K1 and SW_K2, found in two sperm whales sampled in the Kerguelen and SW_C, found in one sperm whale in Crozet. SW_K1 matched the haplotype 10 defined by [45] found off South Australia and Victoria, and SW_K2 corresponded to the haplotype GG [36], exclusively found in the Indian Ocean in the Seychelles. SW_C corresponds to haplotype B [36, 45], found in Australia (Figure 2).

Males from Kerguelen/Crozet shared no first-degree relations with the Irene’s group and had fewer second-degree relationships (n=9, among which only two are found with immatures of the Irène’s group) than Mauritian males (Figure 3). However, some of these males shared strong second-degree relationships with members of the Irène’s group (for example Mystère and Ker 2007, *r*=0.38). Among all adult males sampled off Mauritius or in the south of the Indian Ocean, 24 second-degree relationships were identified (Figure 3, Table S5).

### Average relationship coefficients

During this study, 22 sperm whales (21 adult males and 1 immature female) were added to the 27 already analysed in [21]. The 49 sperm whales in total included in this study were the 25 members of the Irène’s social group, 2 members of another social group, “the Reshna group”, one unidentified female, 13 adult males sampled off Mauritius, and the 8 adult males sampled in Crozet/Kerguelen (the complete list is given in Table S2). The mean relatedness of these different samples and of different combinations were calculated (Figure 4, Table S6). Across all the included individuals (in Mauritius and in Crozet/Kerguelen), we calculated an average *r*=0.046, similar to that calculated between all adult males (*r*=0.044; Table S6). As expected, members of the mostly matrilineal Irène’s group had a higher average pairwise *r* (*r*=0.065, Figure 4).

**Figure 4:**
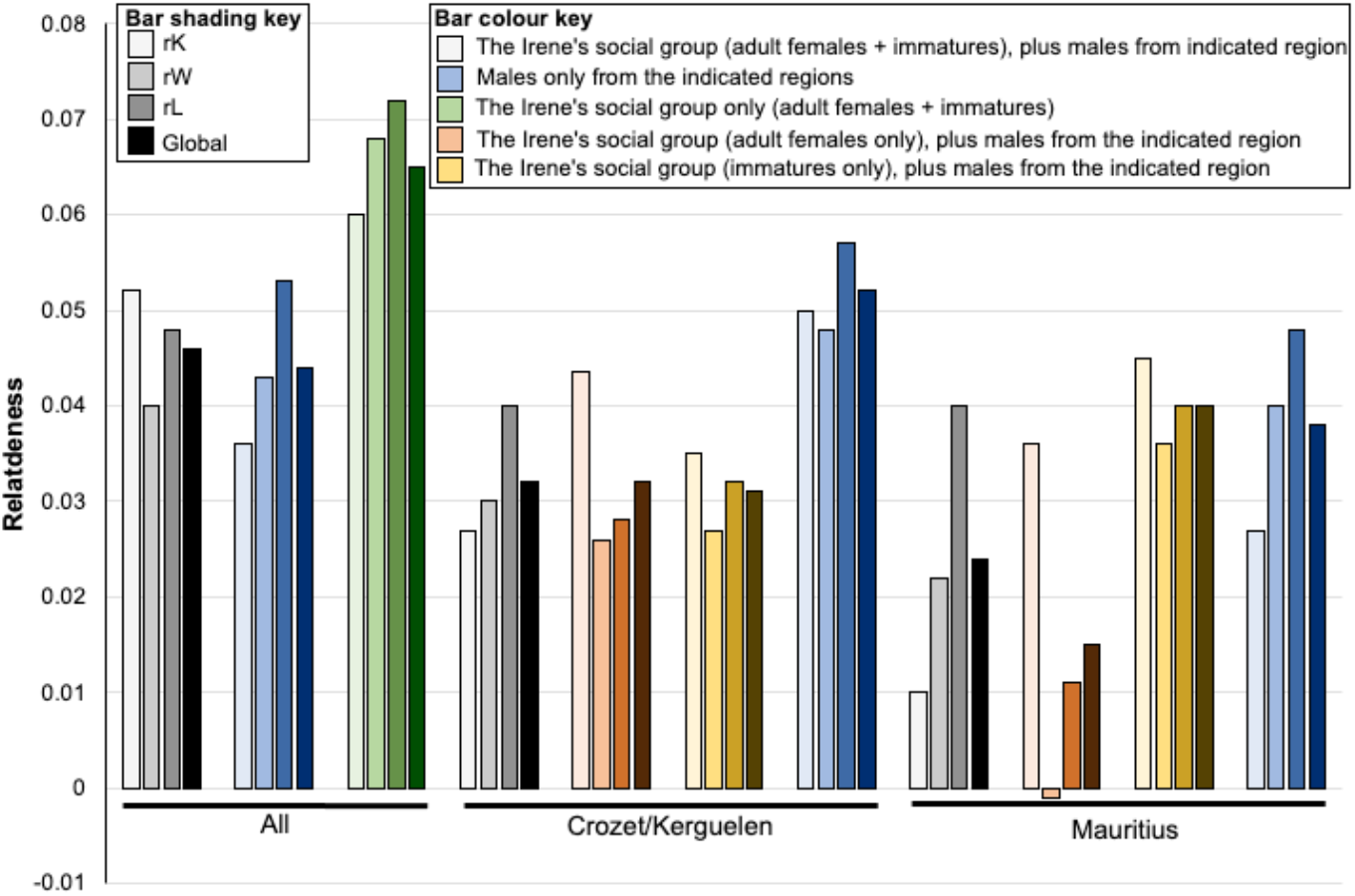
Differences of average relatedness coefficients in groups and subgroups. Relatedness coefficients *r_K_* (Kalinowski *et al*. 2006), *r_W_* (Wang 2002) and *r_L_* (Li *et al*. 1993) were calculated through *ML Relate* and through *Relate. R_Global_ is the average value of the three coefficients (r_K_, r_W_* and *r_L_). The four relatedness estimators are first represented for all individuals, for all adult males and for all members of the Irène’s group. Note that the values of the Irène’s group are higher (shown in green).* *Different combinations of individuals were then formed, and the relatedness coefficients calculated. The partitioning of the Irène’s group between adult females and immatures had a strong impact of the r calculated with adult males sampled off Mauritius, but not with those from Crozet/Kerguelen.*

Average relatedness values were higher among males sampled in Crozet/Kerguelen (*r* = 0.052) than between these males and members of the Irène’s group, whether the Irène group was partitioned into adult females only, immatures only, or the entire group (*r* = 0.031-0.032, Figure 4 and Table S6)

In contrast, the partitioning of the Irène group had an impact on the relatedness values in comparison to males sampled in Mauritius. The average relatedness of the Mauritius males to adult females only from the Irène group was lower (*r* = 0.015) than the relatedness between the immatures and the males (*r* = 0.040, Figure 4 and Table S6).

## Discussion

Currently, our knowledge of behaviour, ecology and genetic diversity of emblematic marine megafauna still suffers from holes. An outstanding example concerns male sperm whales, the “largest toothed creature on Earth” [9]. Sperm whales are steeped in our culture, from the star of one of the most-read novels [46] to the use of their spermaceti oil during the industrial revolution (e.g. [47]). But social and breeding behaviours of male sperm whales remain largely unclear, especially in terms of geographical and social fidelity. Here, we studied sperm whales off Mauritius under the auspices of the Maubydick project [21, 40] and off Crozet/Kerguelen [19, 43, 44]. This allowed us to document the presence of different males visiting the focal mostly matrilineal sperm whale social unit, the Irène group, to identify several recaptures of males with the Irène group over years, to decipher some paternal kinships as well as to capture a diagram of kin relationships at a larger geographic scale. Based on this, we infer that adult males can show social and geographical fidelities to breeding and feeding areas within the Indian Ocean.

### Our study evidenced no natal philopatry of the male sperm whales for the Irène social group

Natal philopatry can be defined as fidelity to birthplace and has been evidenced in different species of marine mammals (e.g. [48, 49]). Among the 13 adult males sampled in Mauritian waters, 12 did not share the SW_M1 haplotype characteristic of the Irène social group [21], and can therefore not have been born in this group. Only one, Herman, had the SW_M1 MCR haplotype, but mitogenome sequencing revealed seven mutations between Herman’s and the predominant Irène’s group mitogenome (Justine Girardet, Agnès Dettaï & Jean-Luc Jung, unpublished). Nuclear DNA analysis is consistent with this statement: the lowest average *r* calculated for any combination of individuals in our study, was between the adult female members of the Irène’s group and males sampled off Mauritius (Figure 4, Table S6).

### Over-years recaptures of different males in the Irène group and estimation of male social fidelity

In contrast to the lack of natal philopatry of adult males demonstrated by our analyses, our study highlights seven examples (three confirmed by resightings over multiple years, and four correlated to gametic recaptures) of males coming back several times to the same area and to the same social unit to breed. These are strong indications that adult male sperm whales may show social fidelity to particular female-dominated social groups, not based on kin relationships with adult females in the group, and that, in turn, they must be well known by the members of these female-dominated social groups.

Nuclear DNA analysis revealed two father-offspring relationships between adult males sampled off Mauritius and immature members of the Irène’s group. One paternity has been attributed to Jonas (father of Daren born in 2018), and one to Noé (father of Lana born in 2019), both sampled in 2018 (figure 4 and Table S4). These “gametic” recaptures [50] proved that some of the males observed in Mauritian waters are reproductive. This reproductive status is supported by the value of the average *r* calculated between males from Mauritius and members of the Irène group, which is nearly tripled if immatures of the Irène’s group alone are considered as compared to adult females of the group (Figure 4, Table S6). The presence of Jonas in the Irène’s group was highlighted over at least three different years (1 year, in 2017, for mating as proved by the “gametic” recapture, and two years of observation, in 2018 and 2019). In addition, nuclear DNA analysis revealed 4 potential full sibling relations. Two are detected between immatures (Alexander and Zoé born in 2019 and 2013, Chesna and Tache Blanche born in 2018 and 2011). The other two are between adult females (Adélie and Emy, Mystère and Irène) whose years of birth are unknown. As twins in sperm whale are very rare [51], it can be assumed that they were not born the same year. Thus, the fathers of each of these four pairs came back at least in two different years to the same group – and to the same specific receptive female – to mate. The father of Chesna and Tache blanche could in addition be the father of Eliot, supposed half-brother of Tache Blanche (Figure 3). Despite these gametic recaptures being based on relatedness estimate calculations, and therefore subject to uncertainties, these findings provide powerful evidence in support of enduring relationships between adult males and specific female-dominated social groups.

It is of note that the three males recaptured between years were seen at the same period of the year (Hugues in October 2013 and October-November 2019, Navin in July 2015 and June 2018, Jonas in July 2018 and May-June 2019). This could indicate either a certain degree of seasonality specific to each individual, or, if they are visiting different female-dominated social groups, a difference in the order that each social group is visited between males. The case of Jonas stands out: Jonas was observed in 2018 and 2019, he is the likely father of Daren, born in 2018, and maybe triggered the gathering of tens of females and immatures in 2018. Jonas has therefore a marked and repeated social fidelity for the Irène’s group, and is in turn well known to the group members. As suggested by Gero et al. [33], spectacular gatherings could also support the hypothesis that females play a role in mating choice.

### Social interactions between adult males and Irène’s group members

Male sperm whales were present in the Irène’s social group most of the year with a peak of occurrence in April and May during the austral autumn, which could represent the breeding season. Labadie et al. [19] and Janc et al. [44] highlighted a seasonality in occurrence of sperm whales in the high latitude feeding area of the Indian Ocean, with increased sightings in spring and summer. However, observations in Mauritius are only conducted daily from February to May, thus the number of males identified in each month could be biased in other months by lower observation effort, therefore reproduction throughout the year cannot be excluded. Residency of males off Mauritius appears to be on the scale of a few days to few weeks with an average stay (8.25d), twice as high as that previously reported off Dominica, for example (3.76d) [33]. Recurrent interactions between adult males and members of the social unit have been observed, confirming previous observations (e.g. [16, 33]). Limited interactions between adult males and adult females and/or immatures have already been reported, for example in Northern Chile and off Dominica [32, 33]. Here, the males identified were often observed in proximity (i.e. less than 100m) of members of the Irène’s social group and several types of interactions (e.g., physical contacts, vocal interactions) were recorded with both adult females and immatures. The exceptional gathering of tens of individuals, - which probably represent a substantial proportion of the local population -, after the arrival of two adult males in the Mauritian waters (MMCO, Field report of July 18 2018) seems to not be restricted to the Indian Ocean: Gero et al. [33] observed a similar aggregation of several tens of individuals near an adult male in the Atlantic. Some males appear therefore to be well known to particular stable social groups. This assumption is reinforced by the numerous interactions observed between adult males and females, and by the several full sibling relationships identified.

### Population genetics and geographical philopatry of male sperm whales in the Indian Ocean

While all members of the Irène’s group except one harboured the same MCR haplotype [21], adult male sperm whales showed a mtDNA diversity in the same range of what was calculated by Alexander et al. [36] for the broader Indian Ocean (Haplotype diversities around *H*=0.8, nucleotide diversities around π=0.0028). The haplotypes identified in this study near Mauritius and matching to Alexander et al. [36] haplotypes all corresponded to minor and major haplotypes of the Indian Ocean. In Crozet/Kerguelen, mtDNA haplotypes suggest a widespread geographic origin of adult male sperm whales: they match to North Indian Ocean haplotypes identified from the west to the east of the Ocean (Figure 2). Even though we sampled only limited numbers of male sperm whales, tests of differentiation based on mtDNA detected some levels of genetic differentiation between Mauritius and Crozet/Kerguelen (*Φ_ST_* and *F_ST_* significant or nearly so), which reflect divergent distribution of mtDNA haplotypes between the two sites, although a high number of second- and third-degree relationships were found between males sampled in the two areas.

While the mtDNA results likely reflect the widespread origin of males at specific geographic locations, nuDNA polymorphisms support male-mediated gene flow at large scales, and highlight the reproductive status of males sampled off Mauritius. Adult males of both areas (Crozet/Kerguelen vs Mauritius) show equivalent numbers of second-degree relations with adult females of the Irène group (seven second-degree relations for the eight males sampled in Crozet/Kerguelen and 11 second-degree relation for the 13 males sampled in Mauritius (Figure 3, Table 1). But many more second-degree relations are found between immatures of the Irène Group and males of Mauritius (n=9) than with those sampled in Crozet/Kerguelen (only two second-degree relations).

The average relatedness *r* calculation revealed similar patterns: between males sampled in Crozet/Kerguelen and members of the Irène’s group, the average *r* is similar when subsetting to adults or immatures of the Irène’s group. Therefore, males sampled in Crozet/Kerguelen do not appear to breed preferentially with the Irène social group. This situation is strongly contrasting with the pattern observed for males sampled in Mauritius, where their role of as paternal relatives was demonstrated by a three times higher average relatedness with immatures than with adult females (Figure 4).

### New insights into adult male sperm whale diversity in the Indian Ocean

Male recaptures and social interactions between males and members of social groups have already been observed and suggest some levels of male social fidelity in breeding areas in the Pacific [38] and in the West Indies [33]. Here, we confirm and extend these observations in the Indian Ocean. The level of this male social fidelity (e.g., for social units, for vocal clans, defined in [20]) is still to be evaluated.

Our results suggest that this fidelity is not due to natal social philopatry, i.e. fidelity for the social group of birth. It appears this behaviour is exclusive to female sperm whales. Therefore, males must acquire their fidelity for places and groups other than that of their birth and based on the diversity of mtDNA haplotypes observed in males, this might occur across large geographical scales.

The high mtDNA diversity found in male sperm whales (as compared to the almost complete absence of diversity found in the group of Irène) is likely to reflect disparities in their respective birth places. Alexander et al. [36] found that, in the Indian Ocean, 44.4% of the variance in mtDNA frequencies was explained by regions, and 12.3% by social groups. If the mostly matrilineal nature of the Irène’s group [21] is a more or less general rule for sperm whale social units in the Indian Ocean, the geographical patterns of mtDNA distributions found by Alexander et al. [36] may well correspond to discrete regional partitions of social units, more than to different proportions of mtDNA haplotypes in different populations, found for instance in humpback whales (e.g. [48, 52]. This would be explained by the strong natal social philopatry of females (more than by a natal geographical philopatry). Interestingly, the situation could well be different in the Pacific, where sperm whale social groups could be of larger size and aggregate more often [11], and where partitioning of variance in mtDNA has been explained by social groups and not by regional differences [36].

The number of adult male sperm whales sampled off Mauritius is relatively low (n=13), but it is nevertheless notable that their mtDNA haplotypes are frequent in different regions of the Indian Ocean neighbouring Mauritius. In contrast, sperm whales sampled in the Crozet/Kerguelen (n=8) have haplotypes found in a much broader area covering all the north of the Indian Ocean, from west to east (this study, [36, 45]). This is reflected by significant or near so *Φ_ST_* and *F_ST_* values between Mauritian and Crozet/Kerguelen males. Mesnick et al. [34] suggested that, in the North Pacific, male sperm whales from different region mix in feeding grounds and exhibit some degree of geographical philopatry for the region of their birth when breeding. Our results highlight a lack of natal philopatry of male sperm whales at the social unit scale but they could well fit into the Mesnick et al. [34] hypothesis, with a certain degree of philopatry at a larger geographic scale (here, an area corresponding more or less to the north west of the Indian Ocean). As in the North Pacific [34], and still remaining cautious because of the low number of samples in our study, the high latitude feeding areas in the Southern Indian Ocean could host mixed groups of male sperm whales with a widespread geographic origin, larger than in the breeding areas. These observations are in perfect agreement with previous population genetic studies, highlighting a strong female philopatry and male-mediated gene flow [17, 23, 36].

## Conclusions

Our results strongly suggest that a double fidelity of adult male sperm whales for breeding and feeding grounds exists in the Indian Ocean: (*i*) a certain level of male fidelity has been detected in feeding grounds of the Indian Ocean [19]; our results, a same male has been sampled in 2011 and 2017 off Crozet), and (*ii*) our study highlights the existence of a social and geographical fidelity in a sperm whale breeding area of the south west of the Indian Ocean.

Until now, sperm whales were not believed to follow defined migration routes [9], but, at least in the Indian Ocean, as some degree of fidelity is now proved both for breeding and feeding areas, male sperm whales could well take similar routes to migrate on successive years, also supported by the similar time of year distinct males were observed when resighted between years. Estimating the strength of both fidelities as well as long-term satellite tags could help to confirm this hypothesis.

## Material and methods

### Field work off Mauritius and skin sample collection

Field work took place off the western coast of Mauritius (Mascarenes Islands, Indian Ocean) between latitudes 20.465°S 57.334°E and 19.986°S 57.605°E, up to 15 km off the coast [21]. Sea surface and underwater observations have been carried out since 2011, under the auspices of a project called *Maubydick* led by the MMCO (Marine Megafauna Conservation Organization, [21, 40]. Since 2015, fieldwork has been conducted almost daily between February and May, and some sporadic observations made during the rest of the year, except in January.

Sperm whales were identified based on specific morphological characteristics (e.g., marks on caudal and pectoral fins and body marks, described in detail in [40]. An “Identity card” was established for each individual and these used to construct a catalogue of individuals [40]. During underwater observation non-invasive samples from individually identified sperm whales were collected from sloughed skin fragments as described by [21].

### Collection of sperm whale biopsies off the Crozet and Kerguelen Archipelagos

The Crozet and Kerguelen archipelagos (Crozet/Kerguelen), located in the subantarctic waters of the south Indian Ocean (respectively 46°S and 49°S), are part of the French TAAF (*Terres Australes et Antarctiques Françaises*). One sperm whale sample came from a stranded male found on the shore of Kerguelen in 2007. The other samples (n=8) were collected between 2011 and 2018 from fishing vessels targeting Patagonian toothfish (*Dissostichus eleginoides*), a fish species targeted by sperm whales, [43, 53, 54]. One sample was taken from a dead individual entangled on a longline [43] and the others were biopsies collected with a crossbow (Barnett Rhino or Barnett Wildcat), which fired a hollow-tipped biopsy dart with a floatable head [55, 56]. All samples were conserved in absolute ethanol. The sampling of sperm whales at Crozet/Kerguelen was approved by the *Comité de l’Environnement Polaire* and the French Ministry of Research (04040.03).

### Molecular methods and analysis

All molecular analysis followed the same methodology as previously described [21, 57, 58]. Briefly, genomic DNA was extracted from the skin and biopsy samples using the NucleoSpin DNA RapidLyse® kit (Macherey-Nagel, Düren, Germany). DNA concentrations were standardized to 10ng/μL. Several molecular analyses were performed for each sample including molecular sexing [59], sequencing of a 638bp fragment of the mtDNA control region (MCR: amplified with the primers DLP1.5 and DLP8G, [50] and genotyping of 18 microsatellites loci (Table S1).

mtDNA sequences were manually edited and aligned with Geneious Pro v.7.1 (Biomatters Ltd, Auckland, New Zealand). The 638bp long MCR fragment used is the same region used in [21]. This fragment overlapped fully with the data from [60] and partially (602bp in common) with the sequences determined by [36]. It also overlapped fully with the 283bp fragment and partially with the 563bp fragment (514bp in common) determined by [45]. A new dataset that included all these sequences was constructed to allow a large-scale comparison between mitochondrial haplotypes. The numbers of haplotypes, the haplotype diversity (*H*) and the nucleotide diversity (π) were calculated using the program DnaSP, V.5.10.01 [61]. The software Arlequin, V3.5.1.2 [62], was used to calculate *F_ST_* and *Φ_ST_*, fixation index estimators for mitochondrial genomes.

Fragment sizes were determined using the “Microsatellite Plugin” of Geneious Pro v.7.1 (Biomatters Ltd, Auckland, New Zealand). All the molecular analyses were performed in at least two independent experiments, from different samples of a same individual when available, or twice from the same sample following Sarano et al. [21]. Twenty-two individuals sampled at least three times between 2017 and 2020 (Table S2) allowed us to estimate the microsatellite-genotyping errors linked to possible poor-quality DNA extracts. We calculated an overall error rate of 2.1% per allele (52 alleles incorrect among the 2432 scored) with this error rate then used in kinship analyses.

### Definition of individual specific genotypes

The procedure of anonymization of the samples described in [21] was also applied to all the samples of this new study to confirm the correspondence between field-identification of individuals (here 13 adult males and an immature female, Chesna sampled only in 2020) and genetic individuals, identified by matching genotypes in the laboratory. Briefly, when collected in the field, each skin sample was assigned to one of the individuals identified and then anonymized with an alphanumeric code. To confirm the validity of the field identifications of skin samples, all the steps of the genetic analyses were performed with anonymized skin samples: samples taken from the same individual were confirmed based on similar genotypes using the Identity Analysis function in CERVUS [63] as described in [21]. Genetic individuals and their corresponding samples are listed in Table S2.

### Kinship analysis

Kinship analyses were performed on the complete dataset (with duplicate samples removed), that is adult females and immatures previously analysed [21] with the newly sampled Chesna (sampled in 2020, Table S2), and all the males sampled in Mauritian waters (n=13) and in Crozet/Kerguelen (n=8). Kinship analysis followed the same methodology as described in [21]. Briefly, we first used different estimators to calculate the relatedness coefficient *r* between all the genotyped individuals using the R package *Related* [64] and the software ML relate [65]. *Related* was used to determine that the *r* estimators W [66] and L&L [67] had the highest correlation between observed and expected relatedness values and were thus selected to calculate the relatedness coefficients. ML relate [65] was used to calculate a relatedness coefficient based on the probabilities of sharing alleles identical by descent, and to assign the most probable familial relationships (among parent–offspring (PO), full sibling (FS), half-sibling (HS), unrelated (U)) to each dyad.

The software Cervus 3.0.7 [63] was also used to assign likely kinships. Based on the combined results of these analysis, all probable first- and second-degree kin relationships [68] were listed. The consistency between familial relationships hypothesized by ML relate and *r* coefficient calculations was analysed for each dyad (see also [21] for a more detailed explanation about this procedure).

## Declarations

### Ethics approval and consent to participate

Permission to conduct the Maubydick project, including the taking of sloughed skin fragments, was granted by the Department for Continental Shelf, Maritime Zones Administration and Exploration of the Mauritius Prime Minister Office, on the 21 February 2017. Skin samples were sent to Brest (France) under the CITES agreement FR1702900025-I. The sampling of sperm whales at Crozet/Kerguelen was approved by the *Comité de l’Environnement Polaire* and the French Ministry of Research (04040.03).

### Consent for publication

Not applicable.

### Availability of data and materials

The datasets supporting the conclusions of this article are included within the article and its additional files. Mitochondrial DNA sequences have been deposited into the Genbank, references MK907146-MK907148, MK907159, MK907163, MK907172, MW854724-MW854731 and MW929445-MW929452.

### Competing interests

The authors declare that they have no competing interests

### Funding

The work in the sub-Antarctic region was supported by the French Polar Institute (IPEV program ORNITHOECO-109). The Lush Foundation partly supported the genetics lab work costs.

## Acknowledgements

We thank Navin Boodhonee and the Blue Water Diving Center (Trou aux Biches, Mauritius) for their all years round field work. A special thanks to Carole Decker (Brest, France) for her help with the lab experiments, and to Marine Pensec and Cédric Le Maréchal (Laboratoire de génétique moléculaire et d’histocompatibilité, CHRU, Brest France) for their help with microsatellite analysis. We are very grateful to Marion Sarano (https://marionsarano.fr/graphiste-freelance/) who made the drawing of the figure 2.

Mauritian public authorities greatly helped the Maubydick project, in particular the Mauritian Prime Minister Office, the Marine Continental Shelf Exploration and Administration (MCSEA; Dr Réza Badal and his team), the Albion Fisheries Research Center (AFRC; Chief Scientific officer Mr Satish Kadhun), the Mauritius Film Development Corporation (MFDC; Mr. Sachin Jootun et Miss Eliana Timol), and the Tourism Authority (TA; Miss Khoudijah Boodoo, ex-Director).

## Authors’ contribution

Both FS and JLJ designed the study. FS, JG, GR, PT, CG, AA and JLJ contributed variously to the conception of the project. FS, VS, RH, AP, GR, PT, CG and HV performed the field experiments and identification of individual sperm whales. JG, JLJ, and AMGS conducted the genetic analysis (laboratory procedures). JG and JLJ analysed and interpreted the genetic data JLJ and JG wrote the manuscript. PT, AA, FS, VSS, BM AMGC, GR, CG, HV and OA critically revised the manuscript.

## Notes

### Competing Interest Statement

The authors have declared no competing interest.

