## Supplementary information for "Long distance runners in the marine realm: New insights into genetic diversity, kin relationships and social fidelity of Indian Ocean male sperm whales"

| Loci | nA | Range (bp) | Repeat type | Reference | Hybridation Temperature | Number of cycles | Dye label |
|---|---|---|---|---|---|---|---|
| Pp HO110 | 11 | 130-150 | Di | Rosel et al. 1999 | 50 | 40 | 6-Fam |
| Pp HO130 | 12 | 126-156 | Di | Rosel et al. 1999 | 55 | 40 | Hex |
| Pp HO102 | 13 | 173-199 | Di | Rosel et al. 1999 | 55 | 40 | Hex |
| Pp HO131 | 4 | 110-120 | Di | Rosel et al. 1999 | 55 | 40 | 6-Fam |
| PPHO133 | 6 | 190-200 | Di | Rosel et al. 1999 | 55 | 40 | 6-Fam |
| PPHO104 | 1 | 132 | Di | Rosel et al. 1999 | 57 | 40 | Hex |
| Ev37 | 20 | 207-249 | Di | Valsecchi & Amos (1996) | 55 | 35 | 6-Fam |
| GT211 | 9 | 186-212 | Di | Berube et al. (2000) | 55 | 35 | 6-Fam |
| GT023 | 6 | 80-90 | Di | Berube et al. (2000) | 55 | 35 | Hex |
| GT575 | 6 | 131-145 | Di | Berube et al. (2000) | 55 | 35 | Dragonfly |
| GATA417 | 4 | 175-191 | Tetra | Palsboll et al. (1997) | 55 | 35 | 6-Fam |
| GATA028 | 5 | 117-137 | Tetra | Palsboll et al. (1997) | 55 | 35 | Dragonfly |
| 199_200 | 1 | 108 | Di | Schlötterer et al. 1991 | 55 | 35 | Hex |
| 417_418 | 3 | 193-197 | Di | Schlötterer et al. 1991 | 55 | 35 | Hex |
| EV1 | 9 | 128-148 | Di | Valsecchi & Amos (1996) | 55 | 35 | Hex |
| EV94 | 11 | 205-225 | Di | Valsecchi & Amos (1996) | 55 | 35 | Hex |
| FCB1 | 10 | 117-141 | Di | Buchanan et al. 1996 | 55 | 35 | 6-Fam |
| FCB17 | 18 | 137-185 | Di | Buchanan et al. 1996 | 55 | 35 | 6-Fam |

**Table S1: PCR conditions, number of alleles (nA) and ranges of allele sizes for each locus.** Microsatellite loci were amplified independently in 20$\mu$L reaction mixes consisting of 10ng of genomic DNA, 1x of *HotStart Taq Mix* (Eurobio®) with 1x ThermoStar® Taq polymerase, 3mM $MgCl_2$, 1mM of dNTPs, 10pmol of each primer (except all PPHO loci, amplified as described in Alfonsi et al. 2012). After an initial denaturing step of 10min at 94°C, N amplification cycles consisting each of three steps (denaturation for 30s at 94°C, annealing for 30s at the locus specific temperature and extension for 1min at 72°C) were conducted. A final extension of 30min at 72°C ended the reaction.

| A. Sperm whales sampled in the Mauritius Island | | | | | |
|---|---|---|---|---|---|
| | Age/sex | 2017 | 2018 | 2019 | 2020 |
| ADELIE | AF | 2017_2B | - | 2019_22 | - |
| AIKO | AF | 2017_03A, 2017_03B, 2017_04B, 2017_05A, 2017_05B, 2017_06B, 2017_07B, 2017_32B | - | 2019_01A | - |
| ALEXANDER | YM | - | - | 2019_12A, 2019_16A | 2020_05 |
| ALI | YM | - | - | 2019_09A, 2019_11A | - |
| AMAN | AM | - | 2018_46A | - | - |
| ANJHIN | AM | 2017_30B | - | - | - |
| ARTHUR | YM | 2017_10B | 2018_10B, 2018_15B, 2018_14B, 2018_36A | - | - |
| CAROLINE | AF | 2017_29B | 2018_18A, 2018_27A | - | - |
| CHESNA | YF | - | - | - | 2020_06 |
| CLAIRE | AF | 2017_25B | 2018_35B | - | - |
| DAREN | YM | - | - | 2019_10A, 2019_09A, 2019_23 | - |
| DELPHINE | AF | 2017_01B, 2017_14B, 2017_20B, 2017_20C, 2017_21B | 2018_08A, 2018_19A, 2018_39A | - | - |
| DOS CALLEUX | AF | 2017_24B | 2018_21A, 2018_29A | - | - |
| ELIOT | YM | 2017_13B | 2018-28A, 2018_36A, 2018_38A | 2019_13A | - |
| EMY | AF | 2017_28B, 2017_33B | 2018_53B, 2018_32A | - | - |
| GERMINE | AF | 2017_23B, 2017_27B | 2018_13B | 2019_18A | - |
| HERMAN | AM | - | - | 2019_20, 2019_21 | - |
| HUGUES | AM | - | - | 2019_37, 2019_38 | - |
| IRENE | AF | 2017_08A, 2017_08B, 2017_09B, 2017_15B, 2017_31B | 2018_01B, 2018_11B, 2018_22A | - | - |
| ISSA | AF | - | 2018_12B, 2018_41A | - | 2020_02, 2020_03 |
| JASON | AM | - | - | 2019_24, 2019_25, 2019_27, 2019_28, 2019_32 | - |
| JEREMIAH | AM | - | - | - | 2020_04, 2020_07, 2020_08 |
| JONAS | AM | - | 2018_45A | 2019_29, 2019_30, 2019_31, 2019_36, | - |
| JOSUAH | AM | - | - | - | 2020_04 |
| LANA | YF | - | - | 2019_08A, 2019_14A, 2019_15A | - |
| LEONARD | AM | - | - | 2019_25, 2019_33, 2019_34, 2019_35 | - |
| LUCY | AF | 2017_26B | 2018_17B, 2018_42A | - | - |
| MINA | AF | 2017_11A, 2017_11B, 2017_12B, 2017_17B | 2018_09B, 2018_20A, 2018_10B | - | - |
| MYSTERE | AF | 2017_16B, 2017_16C | | - | - |
| NOE | AM | - | 2018_34A | - | - |
| Unknown_2017 | AF | 2017_18B | - | - | - |
| REZA | AM | - | - | 2019_03A, 2019_04A, 2019_05A, 2019_06A, 2019_07A, 2019_12A | - |
| ROMEO | YM | - | 2018_30A, 2018_40A | 2019_02A, | 2020_01 |
| Clan_Reshna_1 | AF | - | 2018_03B, 2018_04B, 2018_05B, 2018_06B | - | - |
| Clan_Reshna_2 | AF | - | 2018_07B | - | - |
| TACHE BLANCHE | YM | - | 2018_24A, 2018_26A, 2018_52B, 2018_50B, 2018_33A | - | - |
| TONNERRE | AM | - | - | 2019_38 | - |
| VANESSA | AF | 2017_22B | 2018_25A | - | - |
| VASILILY | AM | - | 2018_43A, 2018_44A | - | - |
| YUKIMI | AF | - | 2018_2B, 2018_39A | 2019_19 | - |

| | | | | | |
|---|---|---|---|---|---|
| ZOE | YF | - | 2018_23A, 2018_31A, 2018_37A, 2018_51B | - | - |

| B. Sperm whales sampled in the Crozet and Kerguelen Archipelagos | |
|---|---|
| Individuals (n=8) | Samples (n=9) |
| PM_Ker_2011_1 | Bio_PM_Ker_2011_1 |
| PM_Ker_2011_2 | Bio_PM_Ker_2011_2 |
| PM_Ker_2011_3 | Bio_PM_Ker_2011_3 |
| PM_Cro_2011_1 | Bio_PM_Cro_2011_1, Bio_PM_Cro_2017_1 |
| PM_Cro_2011_2 | Bio_PM_Cro_2011_2 |
| PM_Cro_2011_3 | Bio_PM_Cro_2011_3 |
| PM_Ker_2016 | Bio_PM_Ker_2016_1 |
| PM_Cro_2018 | Bio_PM_Cro_2018_1 |

**Table S2: Correlation between genetic individuals (skin samples and biopsies sharing a same genotype) and field-identified individuals.**

(**A**) Samples taken off Mauritius. Grey colored boxes indicate samples of adult females and immatures (males and females) already analyzed in Sarano et al. (2021). Thirteen adult males (in white) have been sampled off Mauritius between 2017 and 2020. In the Mauritius Islands, an alphabetic name was given to all the individuals to facilitate individual specific sampling (see Sarano et al 2021). In Crozet and Kerguelen (**B**), individuals are named following the chronological order of the biopsies in the same year. Skin samples represented in red and crossed out are the 7 skin samples attributed to incorrect individuals in the field. Their correct attributions appear in green (see Sarano et al 2021 for explanations).

AF: adult female; AM, adult male; YF, immature female; YM, immature male

| Individual | Number of obs. | Genetic sample | Stay span (in days) | Number of different years of resighting | Dates of observations | | | | | | | |
|---|---|---|---|---|---|---|---|---|---|---|---|---|
| | | | | | 2011 | 2013 | 2015 | 2016 | 2017 | 2018 | 2019 | 2020 |
| Aman | 1 | Yes | 1 | 1 | | | | | | 18 jul | | |
| Anjhin | 10 | Yes | 15 | 1 | | | | | 17, 20, 21, 25, 27, 29 apr<br>01, 02, 04, 05 may | | | |
| Big Frosties | 1 | No | 1 | 1 | | | 30 jul | | | | | |
| Centaure | 1 | No | 1 | 1 | | | | | | | 07 jun | |
| Corto | 1 | No | 1 | 1 | | | | | | | 25 may | |
| Daniel | 1 | No | 1 | 1 | | | | | | | 23 feb | |
| Goliat | 1 | No | 1 | 1 | | | 23 may | | | | | |
| Herman | 4 | Yes | 9 | 1 | | | | | | | 09, 12, 14, 17 apr | |
| Hugues | 4 | Yes | 1-31 | 2 | | 11 oct | | | | | 07, 21 oct<br>06 nov | |
| Jacky | 1 | No | 1 | 1 | | | | | | 03 aug | | |
| Jason | 11 | Yes | 47 | 1 | | | | | | | 23, 26, 29, 30 apr<br>03, 05, 09, 17 may<br>01, 07, 08 jun | |
| Jeremiah | 1 | Yes | 1 | 1 | | | | | | | | 12 mar |
| Jonas | 3 | Yes | 1-30 | 2 | | | | | | 18 jul | 08 may<br>07 jun | |
| Josuah | 1 | Yes | 1 | 1 | | | | | | | | 07 feb |
| Leonard | 11 | Yes | 47 | 1 | | | | | | | 23, 26, 29, 30 apr<br>03, 05, 09, 17 may<br>01, 07, 08 jun | |
| Matsya | 1 | No | 1 | 1 | 19 may | | | | | | | |
| Navin | 2 | No | 1 | 2 | | | 17 jul | | | 16 jun | | |

| | | | | | | | | | | | |
|---|---|---|---|---|---|---|---|---|---|---|---|
| Noe | 2 | Yes | 3 | 1 | | | | | | 16, 18 apr | |
| Reza | 8 | Yes | 15 | 1 | | | | | | | 14, 16, 19, 20, 22, 23, 27, 28 mar |
| Roirené | 1 | No | 1 | 1 | | | | 24 feb | | | |
| Saladin | 1 | No | 1 | 1 | | | | | 14 mar | | |
| Titan | 2 | No | 7 | 1 | | | 11, 17 apr | | | | |
| Titus | 2 | No | 9 | 1 | | | | 21, 29 sep | | | |
| Tonnerre | 1 | Yes | 1 | 1 | | | | | | | 05 dec |
| Ulysse | 1 | No | 1 | 1 | | | | 15 apr | | | |
| Vasilily | 1 | Yes | 1 | 1 | | | | | | 02 jul | |

**Table S3:** Adult male observations near Mauritius over the 9-year study period. The number of days of observation for each male and their date, the interval between the first and last sighting, the number of years a male was observed and the existence of genetic samples are indicated. No observation was recorded in 2012 and 2014.

| | This study | 40 | 189 | 216 | 251 | 266 | 269 | 297 | 609 |
|---|---|---|---|---|---|---|---|---|---|
| | Alexander et al. 2016 | 57 | 206 | 233 | 268 | 283 | 286 | 314 | - |
| Haplotype names | **SW_M1** | **T** | **C** | **T** | **C** | **G** | **A** | **G** | **A** |
| | C.001.002 | • | • | • | • | • | • | • | - |
| | **SW_MC** | **C** | • | • | • | **A** | • | • | • |
| | A.001.001 | C | • | • | • | A | • | • | - |
| | **SW_MCK1** | • | • | • | • | **A** | • | **A** | • |
| | **SW_MCK2** | • | • | • | • | **A** | • | **A** | **G** |
| | N.001.001 | • | • | • | • | A | • | A | - |
| | **SW_M3** | **C** | • | • | • | • | **G** | • | • |
| | KK | C | • | • | • | • | G | • | • |
| | **SW_K2** | • | • | **C** | • | • | • | • | • |
| | GG | • | • | C | • | • | • | • | - |
| | **SW_C** | • | • | • | • | **A** | • | • | • |
| | B.001.001 | • | • | • | • | A | • | • | - |
| | **SW_K1** | • | **T** | • | • | • | • | • | • |
| | **SW_M2** | **C** | • | • | **T** | • | • | • | • |

**Table S4. Name and variable positions of the MCR-haplotypes (in bold).** Correspondence with the haplotype names and numberings of Alexander et al. (2016) is indicated. Position 609 of our dataset has not been sequenced by Alexander et al. (2016). Dots mark similar nucleotides. Hyphens mark undetermined nucleotides.

| | | rw | rl | rk | p | S |
|---|---|---|---|---|---|---|
| **ADELIE** | **ELIOT** | **0,5494** | **0,5866** | **0,53** | **PO** | |
| **ADELIE** | **EMY** | **0,417** | **0,5314** | **0,44** | **FS** | + |
| ADELIE | Clan_Reshna_2 | 0,187 | 0,2007 | 0 | U | |
| AIKO | DAREN | 0,3699 | 0,4487 | 0,38 | HS | |
| AIKO | LUCY | 0,2086 | 0,2007 | 0,19 | HS | |
| AIKO | MINA | 0,1472 | 0,1731 | 0,14 | HS | |
| AIKO | MYSTERE | 0,1292 | 0,2834 | 0,14 | HS | |
| AIKO | Ker_2011_1 | 0,1879 | 0,1731 | 0,2 | HS | |
| ALEXANDER | ALI | 0,4 | 0,3936 | 0,31 | HS | |
| **ALEXANDER** | **CAROLINE** | **0,5745** | **0,5039** | **0,53** | **PO** | * |
| ALEXANDER | ISSA | 0,3437 | 0,3385 | 0,27 | HS | |
| ALEXANDER | JASON | 0,2053 | 0,2282 | 0,11 | U | |
| **ALEXANDER** | **JOSUAH** | **0,2231** | **0,2834** | **0,45** | **HS** | |
| ALEXANDER | LEONARD | 0,4435 | 0,3661 | 0,36 | HS | + |
| ALEXANDER | Unknown_2017 | 0,3111 | 0,2558 | 0,23 | HS | |
| ALEXANDER | VANESSA | 0,0999 | 0,118 | 0,13 | HS | |
| **ALEXANDER** | **ZOE** | **0,4591** | **0,3661** | **0,26** | **HS/FS** | |
| ALEXANDER | Ker_2011_1 | 0,1993 | 0,2007 | 0,14 | HS | |
| ALI | DAREN | 0,2719 | 0,2834 | 0,18 | HS | |
| ALI | DOS_CALLEUX | 0,2945 | 0,3385 | 0,23 | HS | |
| ALI | ISSA | 0,3317 | 0,2558 | 0,09 | U | |
| **ALI** | **MINA** | **0,5688** | **0,5866** | **0,57** | **PO** | * |
| AMAN | Clan_Reshna_2 | 0,3039 | 0,2282 | 0,22 | HS | |
| AMAN | Cro_2011_3 | 0,1761 | 0,0353 | 0,13 | HS | |
| ANJHIN | HUGUES | 0,367 | 0,3109 | 0,25 | HS | |
| ANJHIN | JONAS | 0,1504 | 0,3109 | 0,22 | HS | |
| ANJHIN | REZA | 0,0798 | 0,2007 | 0,13 | HS | |
| ANJHIN | Ker_2011_1 | 0,2544 | 0,2007 | 0,15 | HS | |
| ANJHIN | Ker_2011_2 | 0,369 | 0,2409 | 0,24 | HS | |
| ARTHUR | CHESNA | 0,3051 | 0,2558 | 0,29 | HS | |
| ARTHUR | DELPHINE | 0,2125 | 0,2007 | 0,23 | HS | |
| ARTHUR | EMY | 0,1848 | 0,1731 | 0,21 | HS | |
| **ARTHUR** | **IRENE** | **0,5368** | **0,5039** | **0,51** | **PO** | * |
| ARTHUR | LANA | 0,2555 | 0,2007 | 0,14 | HS | |
| ARTHUR | MYSTERE | 0,2278 | 0,3109 | 0,21 | HS | |
| ARTHUR | REZA | 0,2463 | 0,2558 | 0,25 | HS | |
| CAROLINE | JONAS | 0,174 | 0,2007 | 0,15 | HS | |
| CAROLINE | JOSUAH | 0,1996 | 0,2007 | 0,23 | HS | |
| CAROLINE | LEONARD | 0,2877 | 0,1731 | 0,22 | U | |
| CAROLINE | Unknown_2017 | 0,2232 | 0,2007 | 0,21 | HS | |
| CAROLINE | VANESSA | 0,2262 | 0,118 | 0,15 | HS | |
| **CAROLINE** | **ZOE** | **0,5837** | **0,5314** | **0,58** | **PO** | * |
| **CHESNA** | **DELPHINE** | **0,7007** | **0,7242** | **0,73** | **FS** | * |
| CHESNA | HUGUES | 0,3299 | 0,1727 | 0,22 | HS | |
| CHESNA | IRENE | 0,1901 | 0,2003 | 0,12 | U | |
| **CHESNA** | **TACHE_BLANCHE** | **0,3509** | **0,3658** | **0,42** | **FS** | |

| | | | | | | |
|---|---|---|---|---|---|---|
| CLAIRE | ELIOT | 0,3358 | 0,2007 | 0,14 | HS | |
| CLAIRE | Clan_Reshna_1 | 0,362 | 0,3661 | 0,34 | HS | + |
| CLAIRE | TACHE_BLANCHE | 0,2821 | 0,1456 | 0,12 | HS | |
| DAREN | DOS_CALLEUX | 0,2875 | 0,2834 | 0,25 | HS | |
| DAREN | EMY | 0,2621 | 0,2558 | 0,17 | U | |
| **DAREN** | **JONAS** | **0,4578** | **0,4487** | **0,5** | **PO** | * |
| DAREN | JOSUAH | 0,2156 | 0,2282 | 0,2 | HS | |
| **DAREN** | **LUCY** | **0,5057** | **0,4763** | **0,5** | **PO** | + |
| DAREN | MINA | 0,2634 | 0,2282 | 0,22 | HS | |
| DAREN | MYSTERE | 0,3634 | 0,3661 | 0,27 | HS | |
| DAREN | ROMEO | 0,1939 | 0,1731 | 0,17 | HS | |
| DELPHINE | HUGUES | 0,3578 | 0,2007 | 0,25 | HS | |
| **DELPHINE** | **TACHE_BLANCHE** | **0,5857** | **0,5866** | **0,61** | **PO** | + |
| **DOS_CALLEUX** | **LUCY** | **0,5237** | **0,4763** | **0,5** | **PO** | |
| **DOS_CALLEUX** | **MINA** | **0,5701** | **0,5866** | **0,55** | **PO** | + |
| DOS_CALLEUX | ROMEO | 0,3977 | 0,3385 | 0,33 | HS | |
| DOS_CALLEUX | Unknown_2017 | 0,1936 | 0,1731 | 0,14 | HS | |
| **ELIOT** | **TACHE_BLANCHE** | **0,5283** | **0,5039** | **0,5** | **HS** | + |
| EMY | IRENE | 0,2739 | 0,3385 | 0,24 | HS | |
| EMY | VANESSA | 0,2837 | 0,1731 | 0,15 | HS | |
| EMY | Cro_2011_2 | 0,239 | 0,2282 | 0,17 | HS | |
| GERMINE | HUGUES | 0,0835 | 0,0904 | 0,14 | HS | |
| **GERMINE** | **ISSA** | **0,4464** | **0,3936** | **0,5** | **PO** | * |
| GERMINE | JEREMIAH | 0,1325 | 0,1456 | 0,15 | HS | |
| GERMINE | NOE | 0,2334 | 0,2834 | 0,2 | HS | |
| GERMINE | Ker_2016_1 | 0,1722 | 0,1194 | 0,13 | HS | |
| HERMAN | VANESSA | 0,1366 | 0,0353 | 0,14 | HS | |
| HERMAN | Ker_2011_2 | 0,1762 | 0,1498 | 0,17 | HS | |
| HERMAN | Cro_2011_3 | 0,2404 | 0,1731 | 0,23 | HS | |
| **IRENE** | **LANA** | **0,5378** | **0,4763** | **0,52** | **PO** | |
| **IRENE** | **MYSTERE** | **0,5484** | **0,6141** | **0,54** | **PO** | * |
| IRENE | Ker_2007 | 0,2366 | 0,332 | 0,14 | U | |
| IRENE | Cro_2011_2 | 0,3263 | 0,2834 | 0,13 | HS | |
| ISSA | JASON | 0,2636 | 0,1731 | 0,17 | HS | |
| ISSA | MINA | 0,2043 | 0,2834 | 0,14 | HS | |
| ISSA | Unknown_2017 | 0,2923 | 0,2007 | 0,18 | HS | |
| JASON | Unknown_2017 | 0,2411 | 0,2007 | 0,11 | U | |
| JASON | YUKIMI | 0,1801 | 0,3109 | 0,24 | HS | |
| JEREMIAH | ROMEO | 0,1906 | 0,1731 | 0,18 | HS | |
| JEREMIAH | Ker_2011_1 | 0,1422 | 0,118 | 0,17 | HS | |
| JEREMIAH | Ker_2011_2 | 0,2594 | 0,1801 | 0,16 | HS | |
| JEREMIAH | Cro_2011_2 | 0,2668 | 0,2007 | 0,14 | HS | |
| JONAS | LEONARD | 0,2453 | 0,3385 | 0,21 | U | |
| JONAS | Cro_2011_2 | 0,3008 | 0,3385 | 0,23 | HS | |
| JOSUAH | LEONARD | 0,2411 | 0,3109 | 0,25 | HS | |
| JOSUAH | ROMEO | 0,2046 | 0,1731 | 0,17 | HS | |
| LANA | MYSTERE | 0,2695 | 0,3109 | 0,23 | HS | |
| **LANA** | **NOE** | **0,5224** | **0,559** | **0,54** | **PO** | * |

| | | | | | | |
|---|---|---|---|---|---|---|
| LANA | Cro_2011_2 | 0,4126 | 0,3109 | 0,21 | HS | |
| LEONARD | Clan_Reshna_2 | 0,403 | 0,3661 | 0,3 | HS | |
| LEONARD | VANESSA | 0,0668 | 0,118 | 0,12 | HS | |
| LEONARD | ZOE | 0,4226 | 0,2834 | 0,2 | HS | |
| LUCY | MINA | 0,2114 | 0,2558 | 0,23 | HS | |
| **LUCY** | **ROMEO** | **0,454** | **0,4212** | **0,5** | **PO** | * |
| MINA | ROMEO | 0,1583 | 0,2282 | 0,26 | HS | |
| MYSTERE | Ker_2007 | 0,3756 | 0,4534 | 0,33 | HS | |
| NOE | Ker_2011_3 | 0,397 | 0,332 | 0,28 | HS | |
| REZA | Cro_2018_1 | 0,221 | 0,2282 | 0,24 | HS | |
| Unknown_2017 | ZOE | 0,2756 | 0,2007 | 0,14 | U | |
| Unknown_2017 | Cro_2018_1 | 0,1682 | 0,2282 | 0,14 | HS | |
| Clan_Reshna_1 | Cro_2011_3 | 0,3956 | 0,3385 | 0,35 | HS | + |
| Clan_Reshna_1 | Ker_2016_1 | 0,2944 | 0,2712 | 0,25 | HS | |
| VASILILY | YUKIMI | 0,1935 | 0,2834 | 0,11 | U | |
| VASILILY | Cro_2011_1 | 0,1658 | 0,2007 | 0,18 | HS | |
| VASILILY | Cro_2011_2 | 0,2525 | 0,2007 | 0,12 | U | |
| VASILILY | Ker_2016_1 | 0,2362 | 0,2712 | 0,12 | U | |
| YUKIMI | Ker_2016_1 | 0,2033 | 0,2409 | 0,1 | U | |
| Ker_2011_1 | Ker_2016_1 | 0,2895 | 0,1801 | 0,13 | U | |
| Ker_2011_2 | Cro_2011_2 | 0,2953 | 0,3016 | 0,2 | HS | |
| Ker_2011_2 | Cro_2011_3 | 0,1732 | 0,2105 | 0,21 | HS | |
| Ker_2007 | Cro_2018_1 | 0,3466 | 0,3927 | 0,32 | HS | |
| Cro_2011_1 | Cro_2011_2 | 0,3441 | 0,2282 | 0,16 | HS | |

**Table S5. List of all the first- and second-degree relationships detected between all the individuals.** First (Parent-offspring, PO, and Full siblings, FS) and second (Half siblings, avuncular, grand parents-grandchildrens; all noted HS here) have been deduced
(1) from the calculation of the relatedness coefficient ***r*** using the Kalinowski *et al*. (2006), the Wang (2002) and the Li *et al*. (1993) estimators (respectively $r_K$, $r_W$ and $r_L$),
(2) from the maximum likelihood relationship (*P*.) estimated by ML relate (Kalinowsky et al. 2006) and,
(3) from the parentage analysis performed by CERVUS (*S*: * confidence level 95%, **+**: confidence level 80%, **-**: parent-offspring link undetected by CERVUS).
Twenty-one first degree relationships (PO, n=16 and FS, n=5; all represented in bold) have been detected. Of these relations, two were modified to HS (underlined in the table):
- A FS relation between Alexander and Josuah: impossible as they do not share a mitochondrial haplotype.
- A PO relation between Eliot and Tache Blanche, impossible as Eliot and Tache Blanche are two juveniles born the same year, in 2011, and have different mothers. They most likely share the same father.

| | Average relatedness | | | |
|---|---|---|---|---|
| | **rK** | **rW** | **rL** | **$r_{Global}$** |
| Irene's social group (Adult females and juveniles) | 0.06 | 0.068 | 0.072 | 0.065 |
| All adult males | 0.036 | 0.043 | 0.053 | 0.044 |
| Males sampled in Mauritius | 0.027 | 0.04 | 0.048 | 0.038 |
| Males sampled in Crozet/Kerguelen waters | 0.05 | 0.048 | 0.057 | 0.052 |
| Irene's group with males from Mauritius | 0.01 | 0.022 | 0.04 | 0.024 |
| Irene's group (juveniles only) with males from Mauritius | 0.045 | 0.036 | 0.040 | 0.040 |
| Irene's group (adult female only) with Mauritian males | 0.036 | -0.001 | 0.011 | 0.015 |
| Irene's group with males from Crozet/Kerguelen | 0.027 | 0.03 | 0.04 | 0.032 |
| Irene's group (juveniles only) with males from Crozet/Kerguelen | 0.035 | 0.027 | 0.032 | 0.031 |
| Irene's group (adult female only) with males from Crozet/Kerguelen | 0.044 | 0.026 | 0.028 | 0.032 |
| All the individuals | 0.052 | 0.04 | 0.048 | 0.046 |

**Table S6: Average relatedness coefficients in groups and subgroups**
Relatedness coefficients $r_K$ (Kalinowsky *et al*. 2006), $r_W$ (Wang 2002) and $r_L$ (Li *et al*. 1993) were calculated through *ML Relate* and through *Relate*. $r_{Global}$ *is the average value of the three coefficients (*$r_K$, $r_W$ and $r_L$*)*
